# Crop microbiome responses to pathogen colonisation regulate the host plant defence

**DOI:** 10.1101/2023.02.24.529317

**Authors:** Hongwei Liu, Juntao Wang, Manuel Delgado-Baquerizo, Haiyang Zhang, Jiayu Li, Brajesh Singh

## Abstract

**Aims:** Soil-borne pathogens severely damage the yield and quality of crops worldwide. Plant and soil microbiomes (e.g. in the rhizosphere) intimately interact with the plant, the pathogen and influence outcomes of disease infection. Investigation of how these microbiomes respond to disease infection is critical to develop solutions to control diseases.

**Methods:** Here, we conducted a field experiment and collected healthy and crown rot disease infected (caused by *Fusarium pseudograminearum*, *Fp*) wheat plants. We investigated their microbiomes in different compartments, plant immune responses and interactions with the pathogen (*Fp*) aiming at advancing our knowledge on microbiome-mediated regulation of plant responses to pathogens.

**Results:** We found that *Fp* colonised wheat plants in significant loads, accounting for 11.3% and 60.7% of the fungal communities in the rhizosphere and root endosphere, respectively. However, *Fp* presented with a small fraction of the leaf microbiome, up to 1.2%. Furthermore, *Fp*-infection led to significant changes in the composition of the microbial communities in the rhizosphere and root endosphere while had little impact on leaves. We further found that wheat defence signalling pathways, wheat microbiomes and the pathogen intimately correlated with each other in structural equation modelling. As such, we also identified ecological clusters explained changes in the wheat defence signalling pathways. Lastly, microbial co-occurrence network complexity was higher in *Fp*-infected plants relative to healthy plants, suggesting that *Fp*-infection has potentially induced more microbial interactions in plants.

**Conclusions:** We provide novel evidence that soil-borne diseases significantly disrupt belowground plant microbiomes influencing the responses of plant immunity to pathogens.

## Introduction

Annual crop yield losses caused by plant diseases and pests are estimated at USD$220 billion globally (about 20%∼30% of global harvest), which is a significant challenge to global agricultural production, food security and relevant socio-economic issues (Agrios, 1969; Chakraborty and Newton, 2011). Moreover, the global impacts of soil-borne pathogens is expected to increase as Earth become drier and warmer (Delgado-Baquerizo et al., 2020). Even so, our capacity to mitigate the impacts of soil-borne pathogens on crop production is limited. Many soil-borne fungal pathogens have wide host ranges and survive/grow on organic residues in soils, making it difficult for successive crop cycles to maintain yield (Dean et al., 2012). Also, management of these pathogens in the field such as using agrochemicals is usually ineffective, costly and environmentally unfriendly (Liu et al., 2017). The use of biological tools such as microbial inoculants/crop probiotics has been recommended as an alternative to the use of agrochemicals, e.g. fungicides (Chakraborty and Newton, 2011; Liu et al., 2020b). Increasing evidence suggests that plant microbiomes have a profound influence on the disease triangle that is comprised of the plant, the pathogen and the environment (Liu and He, 2019). Therefore, systematic understandings of the interplay between the microbiome, the pathogen and the plant will enhance our capacity in developing biological tools to control crop pathogens (Liu et al., 2020b). However, unlike cultivar effects on plant microbial heterogeneity, our knowledge about how fungal diseases affect plant microbiomes (diversity, composition and network structure) across different compartments and consequences for the plant growth and health is still limited (Berendsen et al., 2012; Liu et al., 2017; Liu et al., 2020a). This lack of knowledge hampers our ability to understand fundamental ecological interactions between pathogen, microbiomes and plants, and constrain the development of effective disease/farming management strategies.

Plants are associated with many microorganisms in the rhizosphere, phyllosphere and the plant endosphere (Berendsen et al., 2012; Liu et al., 2017; Liu et al., 2020a). These microorganisms, collectively known as plant microbiomes, generally play crucial roles in regulating multiple aspects of the plant growth and stress tolerance (Berendsen et al., 2012; Ritpitakphong et al., 2016; Lu et al., 2018; Liu et al., 2020a). The rhizosphere microbiomes, which are mainly recruited from the soil by the plant, intimately interact with the plant immune system and influence the fitness of the plant. Recent evidence showed that the rhizosphere microbiome directly suppressed the growth of plant pathogens (Chen et al., 2018; Durán et al., 2018). About half of the culturable bacteria in the rhizosphere could suppress the immune system of *Arabidopsis*, which perhaps is an evolutionary mechanism of bacterial symbionts to counteract host immune systems for a better colonisation (Teixeira et al., 2021). Additionally, the composition and diversity of the rhizosphere microbiomes are also determined by the development stage and health status of the plant (Xiong et al., 2021). As such, plants release about 10%- 40% of their fixed carbon into the rhizosphere, which mediates and regulates the rhizosphere microbiome assembly and interactions, especially under stress conditions (Newman, 1985; Bais et al., 2006).

Root endosphere is also considered as a critical interface for plant-microbe interactions, which has been compared as a second layer of plant defence (Dini-Andreote, 2020). Pathogen infection can significantly influence the endosphere microbiomes (Liu et al., 2020b; Trivedi et al., 2020). A recent study demonstrated that sugar beet plants accumulated a bacterial consortium comprised of *Chitinophaga* and *Flavobacterium* in the root endosphere, which could suppress the growth of fungal pathogen *Rhizoctonia solani* in plant tests (Carrión et al., 2019). This suggests that root endosphere microbiomes can be maintained to cope with a particular biotic stress of the plant. Soil- and root-associated microbiomes can transmit to the phyllosphere (e.g., leaf and stem) via xylem and phloem along with other pathways such as soil and air dispersal, insect visitation, and rainfall events (Liu et al., 2017). Recent studies revealed that pathogen infection and/or hormone applications on plant shoots could reshape microbial assemblage in the rhizosphere, with bacteria potentially boosting plant defence capacity being enriched (Carvalhais et al., 2013; Berendsen et al., 2018; Liu et al., 2021). However, it is still unclear if the disease influences the microbiome and induces accumulation of beneficial microbes in the phyllosphere (Liu and Brettell, 2019). Overall, plant microbiomes can modulate plant phenotypes, immunity and defence against pathogens, and respond to stresses in different ways (Liu et al., 2020b). To this end, knowledge gaps still exist, specifically, (i) if there are links between the plant immunity, plant microbiomes and pathogen infection, (ii) whether/how plant microbiomes in different compartments respond to fungal disease infection, and (ii) what are the major microbial taxa enriched/depleted in plants under pathogen attack. Addressing these knowledge gaps is important to identify microbes for improving plant disease resistance and to advance the fundamental disease ecology.

Here, we investigated the impacts of crown rot disease (CR), caused by *Fusarium Pseudograminearum* (*Fp*) on wheat microbiomes. We chose wheat because it is a major global crop and its production is significantly influenced by fungal diseases (Grote et al., 2021). Among these, the crown rot disease is a serious concern, which has recently become more widespread due to the increased adoption of conservation farming (e.g. no-till) in Australia (Yin et al., 2013; Liu et al., 2016b). We hypothesized that (i) a three-way interaction exists between the wheat immune system, the pathogen (*Fp*) and microbiomes. (ii) Pathogenic infection influences wheat microbiomes, which is also compartment-dependent, and the belowground plant microbiomes respond more strongly than leaf microbiomes. And (iii) pathogen loads in wheat differ between plant compartments and correlate with plant disease severity. To test these hypotheses, we collected healthy and *Fp*-infected wheat plants from the WellCamp site in Queensland, Australia. We analysed the *Fp*-induced microbial changes using 16S rRNA and ITS amplicon and shotgun sequencing, and tested correlations between the pathogen *Fp*, wheat microbiomes and defence signalling pathways using structural equation modelling (SEM) and microbial co-occurrence network analyses. By bringing together these analyses, this study aims to unravel how plants influence and modify their microbiomes to mitigate exposure to fungal diseases.

## Materials and method

### Field sampling of wheat plants

Wheat and soil samples were collected from a yield trial (2015) at Wellcamp experimental site (27°33’54.7’S, 151°51’52.0’E), Queensland, Australia (Liu et al., 2021). Stubble residues carrying *Fp* that was inoculated in a CR trial in 2010 have provided inoculums for the CR- infection on wheat (in buffer rows) in this yield trial. Soil chemical properties, histories of farm management (CR trials and fallow periods) and experimental design were previously described (Liu et al., 2021). In this study, we collected leaf, base stem (basal internode), root and the rhizosphere soil from the asymptomatic and symptomatic wheat plants to investigate how wheat microbiomes respond to the natural infection by *Fp* and their relationships with the wheat immunity and CR disease infection. Thirteen weeks after sowing, both the healthy (40) and infected (18) plants were carefully uprooted using a shovel from three different locations of the field and separated into independent individuals. The number of healthy and disease wheat plants reflected the field infection severity by *Fp* (∼30% infection rate from visual rating). For each individual, the top ∼10 cm of two to three leaves were cut, transferred to a 15 mL Falcon tube and frozen in dry ice. The leaf, base stem and root (soil attached) samples were also stored in dry ice and transported to the laboratory within the same day and preserved at -80°C.

### Processing the plant and soil samples, and DNA extractions

The base stem covering brown discoloration of the wheat plant was cut and scored for disease severity (Wildermuth and McNamara, 1994) and genomic DNA (gDNA) was extracted for the quantification of *Fp* abundance in stem. The loosely attached bulk soil were removed from the roots by vigorous shaking, and the rhizosphere soil remained was separated from roots by washing in 25 mL 0.1 M sterile phosphate buffer in a 50 mL centrifuge tube at 200 rpm for 5 min. Roots were then transferred to a new tube. The soil suspension was then centrifuged at 4,000 g for 15 min and the obtained soil pellet was regarded as the rhizosphere soil. The roots in the tube were further fully washed by distilled water, followed by surface sterilisation using 4.0% sodium hypochlorite solution (shaking at 200 rpm for 5 min) to remove microbes on the root surface. The roots were then washed in sterile phosphate buffer for three times, air-dried and grinded in liquid nitrogen for gDNA extraction. Leaf, root and stem genomic gDNA was extracted from about 0.2 g plant materials using a Maxwell^®^ 16LEV Plant DNA Kit on a Maxwell^®^ 16 Instrument (AS2000) according to the manufacturer’s instructions. Soil gDNA was extracted from 0.25 g soil per sample using the PowerSoil^TM^ DNA Isolation kit (MO BIO Laboratories, Carlsbad, CA) using manufacturer’s recommendations. DNA concentrations were determined using a Qubit^TM^ fluorometer with Quant-iT dsDNA HS Assay Kit (Invitrogen). The DNA samples were all stored at -20°C until further analyses.

### Analyses of the plant defence genes involved in the JA and SA signalling pathways

Total RNA isolations from leaf samples were performed using a Maxwell^®^ 16 Total RNA Purification (Promega) kit according to the manufacturer’s recommended protocols. For reverse transcriptase qPCR (qRT-PCR) analyses, 2.5 μg RNA was used for cDNA synthesis using the Tetro cDNA Synthesis kit (Bioline^TM^) as per the recommendations of the kit. Ten defence related genes in the JA and SA signalling pathways were examined to detect changes in plant defence upon *Fp*-infections. The genes included *TaAOS* (*Triticum aestivum* allene oxide synthase), *PR2* (beta-1,3-endoglucanase), *PR3* (*Chi1* gene), *PR4a* (*wheatwin 1-2 gene*), *PR5* (a thaumatin-like protein), *PR10* (a wheat peroxidase), *TaPAL* (phenylalanine ammonia lyase), *Lipase*, *TaNPR1* (nonexpressor of pathogenesis-related genes 1) and *WCI3* (wheat chemically induced gene), which were quantified using the SYBR Green qRT-PCR kit on a ViiA™ 7 sequence detection system (Liu et al., 2016a). The qRT-PCR system, thermal conditions, primer sequences and analysing methods were detailed previously (Liu et al., 2021).

### Quantification of Fp in wheat base stem

*Fp* abundance in wheat base stems was quantified by qPCR analyses using the *Tri5* gene (trichothecene cluster responsible for trichothecene production) of *Fp* (Melloy et al., 2010). Total fungal abundance in stem was measured using ribosomal 18S rRNA gene (Melloy et al., 2010). Wheat actin-binding protein coding sequences were used as the reference gene for the analyses of both the *Fp* and total fungal abundance. qPCR was performed using SYBR Green on the ViiA™ 7 sequence detection system. The qPCR system, thermal conditions and primer sequences were previous described (Liu et al., 2021). Gene amplifications were specific as indicated by melt curve analyses. Biomass of *Fp* was then calculated using the formula below

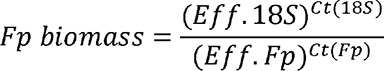

where *Eff* is PCR amplification efficiency calculated by LinRegPCR7.5 (Ramakers et al., 2003).

### Profiling microbial communities in plant and soil samples using high throughput 16S rRNA and ITS amplicon sequencing and metagenomics analyses

We used the 799F-1193R primer set for profiling the root and leaf associated bacterial communities as these primers help to avoid amplification of chloroplasts and other plant-associated DNA sequences (Horton et al., 2014). As the risk of plant contamination is reduced in bulk and rhizosphere soil we used the 926F (Engelbrektson et al., 2010) and 1392R (Peiffer et al., 2013) primer set which are more generally in our lab. Our objective was to compare healthy and *Fp*-infected communities within plant compartments and throughout the paper, we ensured not to make statistical comparisons between inventories generated with different primer sets. FITS7 (Ihrmark et al., 2012) and ITS4 (Innis et al., 2012) were used to amplify fungal communities in all plant and soil samples. Microbial amplification products for plant samples were obtained by running PCR products on gels, where microbial amplicons (∼400 bp) were cut and then cleaned using a Wizard® SV Gel and PCR Clean- Up System (Promega). The plant mitochondrial DNA-derived amplicons (∼800 bp) were discarded. The bacterial communities of the plant and soil samples were sequenced on an Illumina MiSeq as per the manufacturer’s instructions at the University of Queensland, while fungal communities and metagenomics analyses (sequenced using NovaSeq platforms) were conducted using a standard protocol by the Western Sydney Next Generation Sequencing (NGS) facility (Sydney, Australia). Bioinformatic analyses are described in the Supplementary Materials of this study.

### Statistical analyses

Statistical analyses were implemented in R4.0.3 unless otherwise stated. Effects of *Fp-* infection on wheat microbial community composition were investigated using permutational multivariate analysis of variance (PERMANOVA, permutation=9999) and visualised with non-metric multidimensional scaling (NMDS) or principal component analysis (PCA) using the Vegan package (v.2.5-6) (Oksanen et al., 2007). Linear model (Pearson correlation) was performed to examine correlations of the abundance of bacterial and fungal taxa with *Fp* amounts in plants using the package ggpubr (0.1.6) (Kassambara, 2018). To identify marker OTUs that distinguish the healthy and diseased wheat plants, random forest tests were performed using an online tool (https://www.microbiomeanalyst.ca/MicrobiomeAnalyst/home.xhtml) (Chong et al., 2020).

### Structural equation model analyses

Structural equation modelling was used to build a system-level understanding of the effects of *Fp*-infection and plant defence signalling pathways on the variation of the rhizosphere microbiomes. The maximum-likelihood estimation was fitted to the model, and Chi-square and approximate root mean square error were then calculated to evaluate the effectiveness of the model fit. We interpreted a good model fit as one with a non-significant chi-square test (*P* > 0.05), high goodness of fit index (GFI) (> 0.90), low Akaike value (AIC) and root square mean error of approximation (RMSEA) (< 0.05), as previously described (Delgado-Baquerizo et al., 2016). SEM analyses were conducted using the AMOS 22 (IBM, Chicago, IL, USA) software.

### Microbial co-occurrence network analyses

The microbial co-occurrence network analysis that reveals potential interactions between microbial symbionts was used to explore microbial interactions within the wheat microbiomes (Faust and Raes, 2012; Delgado-Baquerizo et al., 2018). In brief, bacterial/fungal OTUs exist across at least three samples were kept for network calculations to reduce spurious correlation that can be caused by rare taxa. To reduce false discovery rate, Benjamini-Hochberg corrections based on 100 bootstraps were conducted and only correlations greater than 0.4 and *P* less than 0.05 were maintained. This cut-off allows the analysis only focusing on taxa that are likely to interact with each other within microbiomes. Correlations were calculated using the SparCC-based (Friedman and Alm, 2012) algorithm Fastspar (Watts et al., 2019). We then identified key ecological clusters/modules comprised of both bacterial and fungal communities in the root environment. The relative abundance of each ecological cluster was computed by standardising the data (z-scores) of the taxa from each module. This allows to exclude any effects of merging data from different microbial groups. Pairwise Spearman’s (q) rank correlations between all taxa (% relative abundance) were used and we exclusively focused on positive correlations because they provide information on species that may respond similarly to disease infection by *Fp*. Only robust Spearman’s correlation coefficient (*R*>0.25 and *P*<0.01) was maintained. All networks were visualised and edited in Gephi, with default network resolution (=2.0) being used to identify ecological clusters (Bastian et al., 2009).

## Results

### A three-way interaction between wheat plant defence, pathogen infection and microbiomes

All the collected wheat plant samples were evaluated with disease severity using qPCR method targeting the *Tri5* gene, and those plants with a gene abundance>1 were scored as disease-infected plants (consistent with visual ratings based on stem discoloration) (Fig.1A). We then investigated *Fp*-induced changes in microbial clusters (network modules) in wheat microbiomes, and their correlations with the pathogen (*Fp*) and plant molecular defence (JA and SA signalling pathways) using SEM analyses and network analyses (Fig.1B, C and D). By doing so, we first built a microbial network across all samples with different ecological clusters (Fig.1B). The abundance of each ecological cluster in the healthy and diseased plants was then calculated and fitted into the SEM to detect direct and indirect interactions between the plant disease, plant defence and microbiomes (Fig.1C and D). Overall, four main ecological clusters were identified, namely C0, C1, C2 and C3, which were comprised of bacterial and fungal taxa from different phyla with different abundances (Fig.1B and C). We found that *Fp*-colonisation in wheat base stems directly contributed to the occurrence of CR- disease symptoms in wheat (*R*=0.71, *P*<0.05). The CR disease infection then significantly contributed to changes in wheat microbiomes (both the bacterial and fungal communities) in the rhizosphere and root endosphere, driven by C1 and C3 (C1, *R*=0.41, 0.05<*P*<0.1; C3, *R*=0.63, *P*<0.05). Importantly, C3 also contributed to changes in the wheat SA signalling pathway (*R*=-0.20, 0.05<*P*<0.1) (Fig.1C), and C3 likely had a higher abundance in the healthy plants relative to diseased plants (Fig.1C). Consistently, the abundance of particular microbial taxa (e.g. OTU_89589 and OTU_41442, *Stenotrophomonas* sp.) in wheat microbiomes significantly correlated with transcript abundances of the defence genes (Table S1), suggesting that wheat microbiomes have a role in regulating plant molecular defences. Lastly, *Fp-*infection significantly contributed to the enhanced gene expression in wheat JA signalling pathway (*R*=0.63, *P*<0.05) (Fig.1D).

**Fig. 1.**
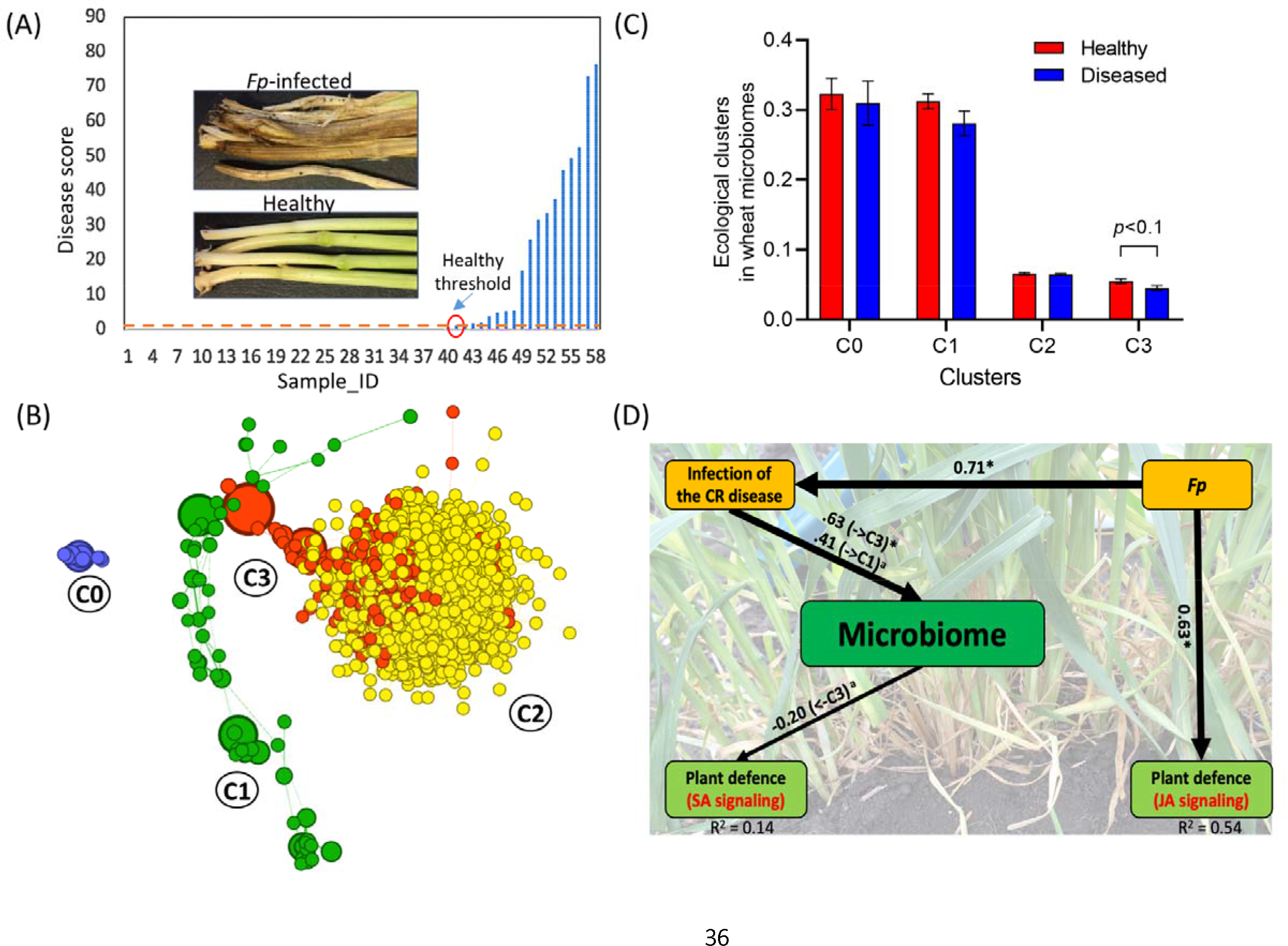
Correlation network analyses of the wheat-associated microbiomes (bacterial and fungal communities) in the rhizosphere and root endosphere. (A) Disease scores for wheat plants obtained from qPCR analyses targeting *Tri5* gene of *Fp*. The inserted pictures showed *Fp*- infected and health wheat stems, which had brown discolorations and normal green colours respectively. The red dash line indicates the threshold for heathy plants (disease score<1.0); (B) Network diagram with nodes coloured according to each of the four main ecological clusters (C0–C3); (C) The relative abundance of each ecological cluster in the healthy and diseased wheat plants (0.05<*P*<0.1); (D) Structural equation modelling (SEM) summarising direct and indirect effects of plant signalling pathways and pathogen infection on wheat microbiomes. Solid lines: positive correlations (significance levels \**P* < 0.05, ^a^ 0.05<*P*<0.1). Numbers around the arrows are indicative of the correlations, and the proportion of explained variance (R^2^) appears alongside each factor in the model. Error bars in B represent standard errors of the mean. Abbreviations: *Fp*, *Fusarium pseudograminearum*; CR, crown rot; SA, Salicylic acid; JA, Jasmonic acid.

**Table 1.** Enriched or depleted bacterial and fungal taxa in wheat microbiomes based on changes in relative abundances before and after Fp ection. Rel-Abun: relative abundance. A significant P value threshold was set at 0.01 to minimize spurious correlations.

| Rhizosphere |  |  |  |  |  |  |  | Root endosphere |  |  |  |  |  |  |  |
| --- | --- | --- | --- | --- | --- | --- | --- | --- | --- | --- | --- | --- | --- | --- | --- |
| Bacteria |  |  |  | Fungi |  |  |  | Bacteria |  |  |  | Fungi |  |  |  |
| Taxa | *R* | *P* | Rel-Abun | Taxa | *R* | *P* | Rel-Abun | Taxa | *R* | *P* | Rel-Abun | Taxa | *R* | *P* | Rel-Abun |
| *Stenotrophomonas*<br>sp. | 0.65 | <0.0001 | 0.08-4.86% | *Fp* | 0.76 | <0.0001 | 0-11.3% | *Stenotrophomonas*<br>sp. | 0.47 | <0.001 | 0.07-11.4% | *Fusarium* sp.<br>(not the<br>pathogen) | -<br>0.39 | <0.001 | 0.35-24% |
| *Pseudomonas* sp. | 0.42 | <0.01 | 0-1.63% | *Sarocladium*<br>*strictum* sp. | 0.39 | <0.01 | 0-2.05% | *Massilia* sp. | -<br>0.38 | <0.01 | 0.25-3.79% | *Fp* | 0.63 | <0.0001 | 0-60.7% |
| *Rhizobacterium*<br>sp. | 0.4 | <0.01 | 0.17-1.04% | *Coprinellus*<br>sp. | 0.45 | <0.001 | 0-2.14% | *Rhodoferrax* sp. | 0.48 | <0.001 | 0-1.01% | *Hydropisphaera*<br>sp. | -<br>0.45 | <0.001 | 0.01-13.6% |
| *Microbacterium*<br>sp. | 0.37 | <0.01 | 0-1.73% |  |  |  |  | *Massilia* sp. | -<br>0.44 | <0.001 | 0.04-1.08% | *Mortierella*<br>*alpina* | -<br>0.36 | <0.01 | 0.1-2.8% |
| *Arthrobacter* sp. | -<br>0.35 | <0.01 | 0.17-1.17% |  |  |  |  |  |  |  |  | *Fusarium* sp. | -<br>0.35 | <0.01 | 0%-<br>2.4% |

### Fp-infection altered microbiomes in the wheat rhizosphere and root endosphere but not in the leaf

By using the 16S rRNA and ITS amplicon sequencing, we found that the bacterial and fungal communities in both the rhizosphere and root endosphere were significantly influenced by the *Fp*-infection (Fig.2C, D, E and F). Such influences on microbiomes were restricted to the community composition, but not the alpha diversity (Table S2). Furthermore, the bacterial community in the rhizosphere (Fig.2D, *P*=0.0002, stress=0.18) responded to the disease more prominently than the root endosphere (Fig.2E, *P*=0.073, stress=0.11). In contrast, bacterial (*P*=0.23, stress=0.16) and fungal (*P*=0.22, stress=0.16) community composition of the leaf was not affected by the *Fp*-infection (Fig.2A and B). The alpha diversity (Observed OTUs, Chao1 and Shannon) of the leaf microbiome was also not influenced (Table S2).

**Fig. 2.**
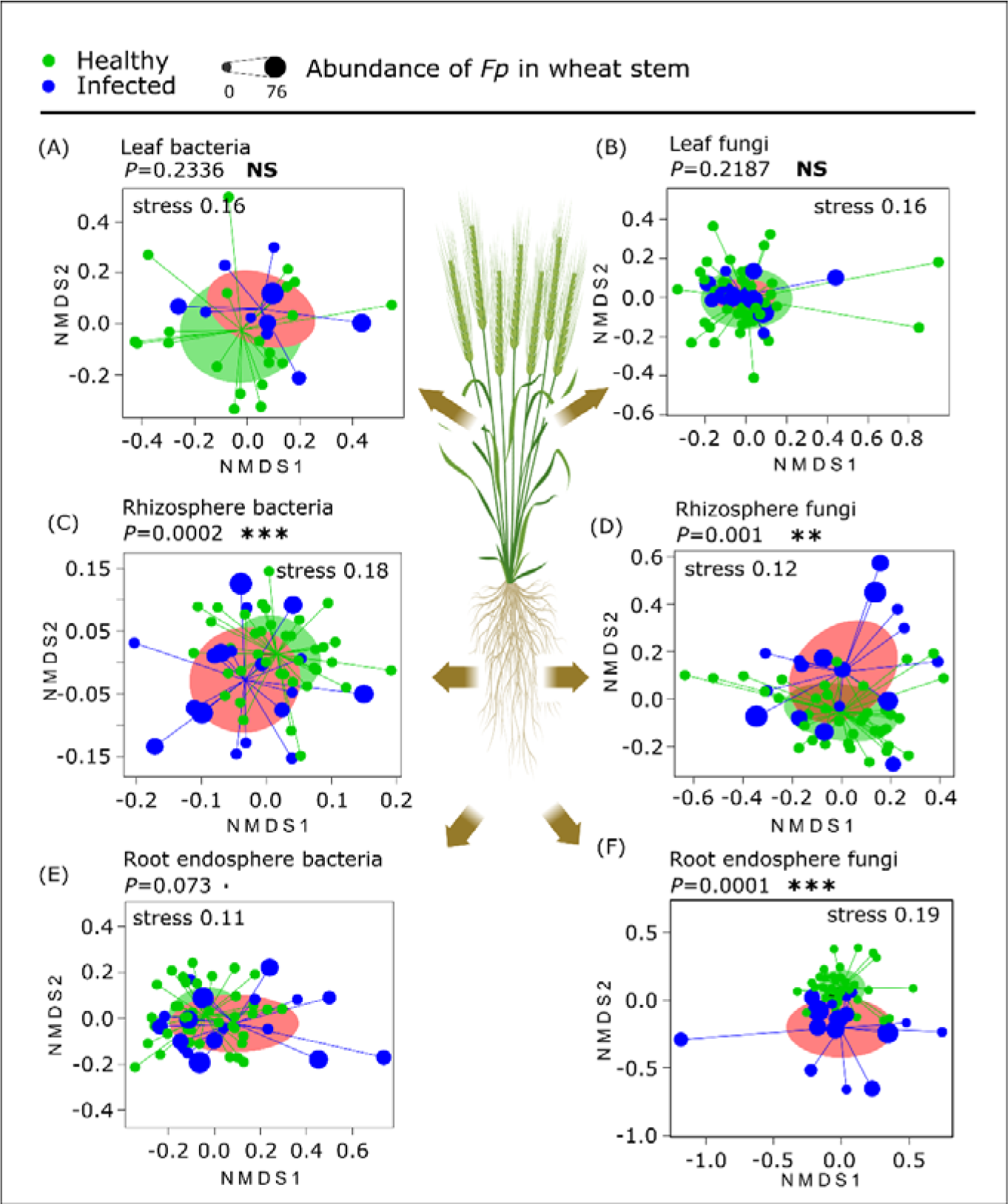
Nonmetric multidimensional scaling ordination summarising impacts of *Fp*-infection on wheat microbiomes in different compartments. (A) Leaf bacterial community, (B) leaf fungal community, (C) rhizosphere bacterial community, (D) rhizosphere fungal community, (E) root endosphere bacterial community, and (F) root endosphere fungal community. Each of the dot in the graphs represents a plant/soil sample, and was scaled to the *Fp* abundance in the wheat base stem (\**P* <0.05, ** *P*<0.01, \*\*\**P*<0.001). NS: non-significant.

### Enrichment/reduction of abundant bacterial taxa in the root and the rhizosphere

At the operational taxonomic units (OTU) level, we detected five abundant bacterial OTUs in the rhizosphere (>1.0%, relative abundance) which had significant positive or negative linear correlations with the *Fp*-abundance in base stem (a direct measurement of CR disease severity). These OTUs included a *Stenotrophomonas* sp. (OTU_89589, *R*=0.65, *P*<0.0001), a *Pseudomonas* sp. (OTU_85258, *R*=0.42, *P*<0.01), a *Rhizobium* sp. (OTU_121632, *R*=0.4, *P*<0.01), a *Microbacterium* sp. (OTU_117739, *R*=0.37, *P*<0.01) and an *Arthrobacter* sp. (OTU_113799, *R*=-0.35, *P*<0.01). For the bacterial community in the root endosphere, those OTUs significantly changed in relative abundances included a *Stenotrophomonas* sp. (OTU_41442, *R*=0.47, *P*<0.01), two *Massilia* spp. (OTU_21226 and OUT_61703, R=-0.38 and -0.44, *P*<0.01 for both cases) and a *Rhodoferax* sp. (OTU_53413, R=0.48, *P*<0.01). Among these, *Massilia* spp. decreased while the *Rhodoferax* sp. increased in relative abundance in wheat microbiomes by the *Fp*-infection (Table 1). Lastly, we did not detect any enriched/depleted bacterial taxa in the wheat leaf.

### A progressive route for Fp-infection and its impact on fungal communities in the rhizosphere, root endosphere and leaf

The composition of what fungal microbiomes significantly differed between compartments (*P*<0.0001, R^2^=0.46), with the leaf fungal microbiome being well separated from those in the root, rhizosphere and bulk soil along the X axis of the principal component analyses (PCA) (Fig.3A). When analysing each of the fungal OTU in the rhizosphere, root endosphere and leaf samples independently (Fig.3A), OTU_13 (ID: f2715a0278527493146542b156252d9d) in the rhizosphere had a strong positive linear correlation with the CR disease severity (measured by qPCR method targeting the *Tri5* gene) in wheat plants (*R*=0.76, *P*<0.0001) (Fig.3B). OTU_13 was identified as a f_Nectriaceae using the UNITE database (https://unite.ut.ee/). When blasted the amplicon sequence (241 bp) in National Center for Biotechnology Information (NCBI), we found it had a 100% nucleotide similarity to a *Fp* strain (ID: MT465499.1) (Fig.S1). These results strongly suggested that OUT_13 was the pathogen (*Fp*) that caused the CR disease on wheat in the field experiment. Its abundances reached to 11.3% and 60.7% in the rhizosphere and root endosphere fungal communities, respectively (Table 1). Also, its relative abundance in the root endosphere was significantly correlated to the *Fp* abundance in wheat base stem (*P*< 0.0001).

**Fig. 3.**
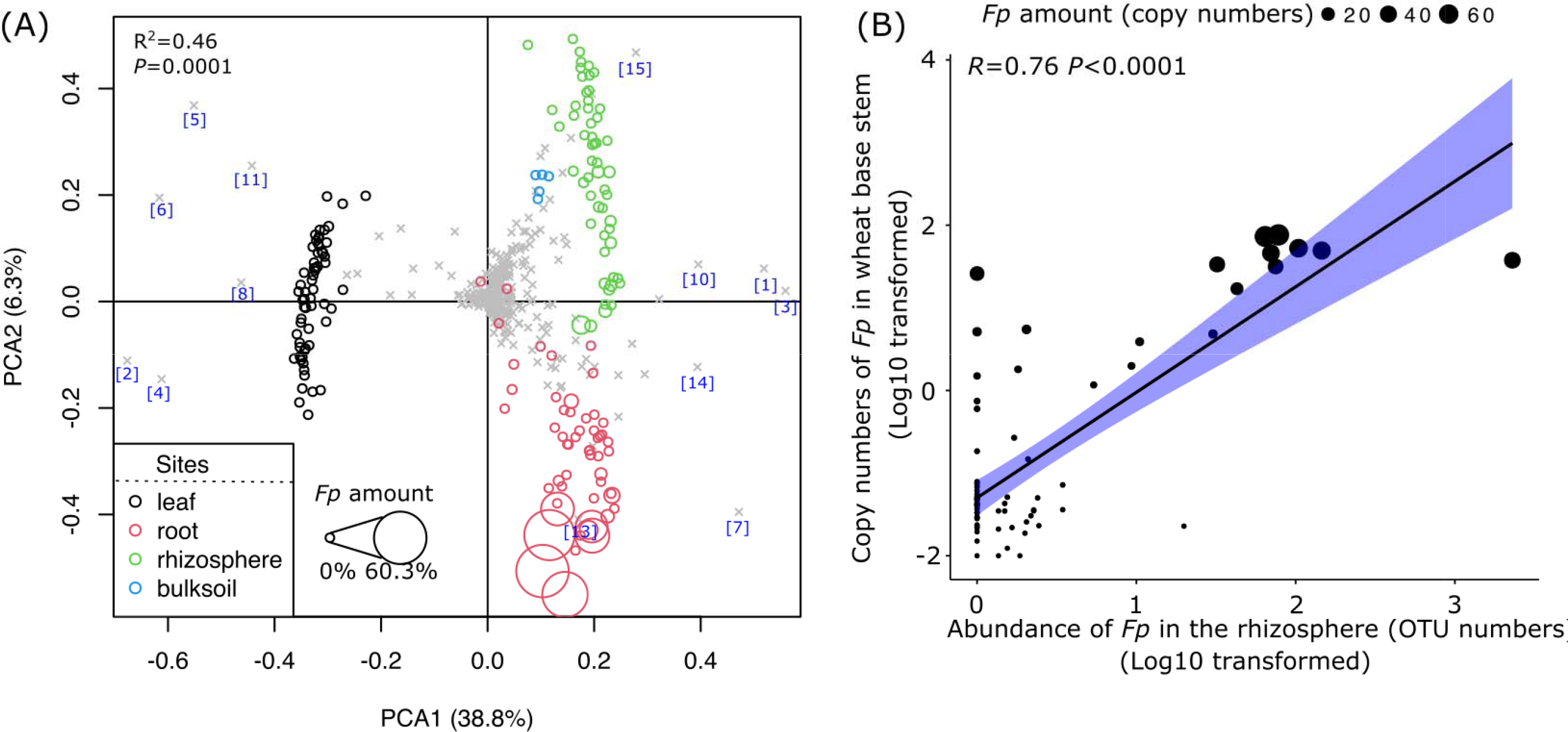
Changes in fungal community structure and pathogen load in different compartments. (A) Principal analysis (PCA) summarising differences in fungal community composition between the bulk soil, rhizosphere, root endosphere and leaf samples. The numbers in brackets in A represent major OTUs driving the separation of fungal communities between different plant compartments and bulk soil. OTU_1: *Fusarium* sp., OTU_2: *Blumeria* sp., OTU_3: *Emericellopsis* sp., OTU_4: *Blumeria* sp., OTU_5: Pleosporales, OTU_6: *Cladosporium* sp., OTU_7: *Olpidium* sp., OTU_8: *Blumeria* sp., OTU_10: *Oidiodendron* sp., OTU_11: Didymellaceae, OTU_13: *Fusarium pseudograminearum* (the fungal pathogen), OTU_14: Hypocreales sp., and OTU_15: *Mortierella* sp. Each of the circle in the graph represents a plant/soil sample, with the size being scaled to the *Fp* amount in wheat base stems. (B) A significant linear correlation between OTU_13 in the rhizosphere (obtained from ITS amplicon sequencing) and *Fp* load in wheat base stem. The size of dots was scaled to the abundance of *Fp* in wheat base stems (*Fp* amount was log10 transformed). The solid line represents the linear regression and grey shaded area represents 95% confidence. OTU: operational taxonomic unit.

Besides OTU_13, two other fungal taxa in the rhizosphere, OTU_2568 (*Sarocladium strictum* sp., *R*=0.39, *P*<0.01) and OTU_949 (*Coprinellus* sp., *R*=0.45, *P*<0.001) also had positive linear correlations with the *Fp*-load in wheat base stem (Table 1). In the root endosphere, OUT_1 (*Fusarium* sp., *R*=-0.39, *P*<0.001, not matching with the pathogen sequences), OUT_14 (*Hydropisphaera* sp., *R*=-0.45, *P*<0.001), OUT_32 (*Mortierella alpina*, *R*=-0.45, *P*<0.01) and OUT_42 (*Fusarium* sp., *R*=-0.35, *P*<0.01, not matching with the pathogen sequences) had negative correlations with the disease severity of the wheat plant. These results suggested that invasion of the wheat plant by *Fp* had significantly interrupted the fungal community composition but not the alpha diversity (Table S2). In the leaf, OTU_13 reached up to 1.2% of the total fungal community, and its relative abundance did not correlate to the disease severity (*R*=0.11, *P*=0.39). We did not detect OTU_13 in the bulk soil.

We then identified key microbial taxa distinguishing the wheat microbiomes of the healthy and diseased wheat plants using random forest tests (Fig.S2). For the rhizosphere bacterial community, the 10 OTUs included a *Stenotrophomonas* sp., a *Saccharothrix* sp. an *Arthrobacter* sp., two *Massilia* spp., a *Sphingopyxis* sp., a Gemmatimonadaceae, a *Lysobacter* sp., and an Oxalobacteraceae (Fig. S2C). Among these, *Stenotrophomonas* sp. was much more effective in distinguishing diseased and healthy plant-associated microbiomes than other taxa. For the fungal community, OTU_13 was the most effective in separating the healthy and diseased wheat microbiomes in the rhizosphere (Fig. S2D).

### Impacts of Fp-infection on co-existence network structure of wheat microbiomes in different compartments

To examine the effects of *Fp*-infection on potential microbial interactions in different wheat compartments, microbial co-existence network analyses were conducted (Fig.4). Consistent with changes in microbial community composition (Fig.2), *Fp-*infection induced changes in microbial networks that were more prominent in the root/rhizosphere environments than the phyllosphere (Fig.4 A, B, C, D, E and F). *Fp*-infection induced a more complex network (in terms of numbers of nodes and edges in networks) for both the bacterial and fungal communities in the wheat rhizosphere and root endosphere than in healthy plants (Fig.4G). The rhizosphere fungal communities had the highest number of nodes and edges, followed by the bacterial communities in the root endosphere and rhizosphere; the leaf-associated microbiomes had the lowest number of nodes and edges for both the bacterial and fungal communities (Fig.4G). The OTUs with high abundances in diseased plants, such as OTU_89589 (*Stenotrophomonas* sp.), OTU_41442 (*Stenotrophomonas* sp.), OTU_121632 (*Rhizobium* sp.), and OTU_85258 (*Pseudomonas* sp.) also had relatively high connections with other taxa in the diseased plants.

**Fig. 4.**
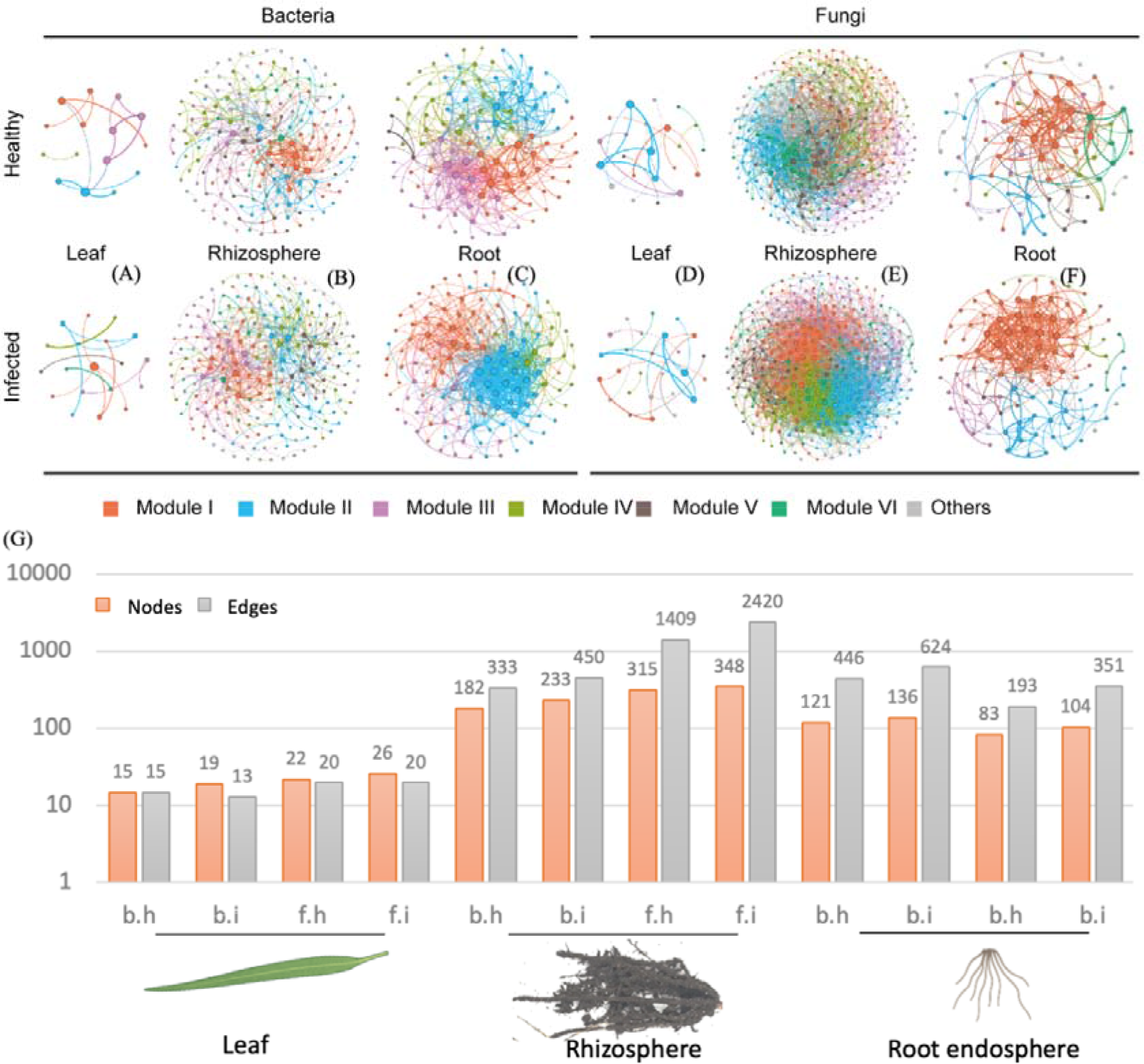
Microbial co-occurrence network analyses of wheat microbiomes. (A-F) Six different major modules were identified, which differentially distributed in the healthy and infected plant microbiomes. (G) Topology of co-occurrence network analyses of the plant/soil microbiomes. Abbreviations: b, bacterial community; f, fungal community; h, healthy wheat plants; and i, infected wheat plants.

### Functional genes and their profile changes in the rhizosphere upon Fp-infection

To detect changes in functional gene abundances and their implications in plant defence and disease resistance/infection, we randomly selected three *Fp*-infected and three healthy wheat rhizosphere soil samples for metagenomic analyses. Consistent with results of 16S rRNA and ITS gene amplicon sequencing, no significant differences were observed for alpha diversity among samples. In terms of beta-diversity of the rhizosphere bacterial community and Genomes (KEGG) Orthology (KO) functional gene composition, we found that the healthy and diseased samples were separated in PCA analyses along the X axis but being only marginally significant for statistics (functional genes, R^2^=0.21, *P*=0.09) (Fig.S3A) and microbial community composition (R^2^=0.30, *P*=0.09) (Fig.S3B). *Fp*-infection induced changes in a diverse range of microbial functions in the rhizosphere including those genes involved in amino acid and carbohydrate metabolisms and signal transductions (Table S3). In total, 30 unique KOs were significantly differed between the disease and healthy wheat rhizosphere soil samples (Table S3). For example, genes encoding glutaryl-CoA dehydrogenase, alpha-trehalase, quinolinate synthase and isocitrate dehydrogenase reduced in relative in the *Fp*-infected wheat plants abundance (by -16.4% to -25.6%), and eleven genes (e.g. those encoding chorismite mutase and sugar phosphate sensor protein UhpC) were enriched in *Fp*-infected plant rhizosphere, but appeared to be in low relative abundance.

## Discussion

Our study investigated the response of wheat microbiomes and immune system to *Fp*- infection on the plant and their correlations with the wheat defence signalling pathways. The findings provide novel evidence for a three-way interaction among the plant defence, the pathogen and plant microbiomes. This knowledge provides novel frameworks for better understanding plant microbiomes and to manipulate them for an improved plant resistance to pathogen attacks. Our results revealed that pathogen (*Fp*) colonisations in the infected plants largely differed between plant compartments and those of the root endosphere and rhizosphere significantly correlated to the CR-disease severity. In contrast, pathogen loads in the wheat leaf of both infected and healthy plants were very low. Furthermore, our study provides strong evidence that *Fp*-infection influences the composition and network structure (complexity and size) of wheat microbiomes in the rhizosphere and root endosphere while *Fp*-infection does not affect the bacterial or fungal communities of the leaf.

### A three-way interaction between wheat defence, microbiomes and pathogen infection

Our finding of a three-way interaction between the wheat defence, microbiomes and the pathogen provides novel evidence of a microbiome role in plant defence and immune responses. The plant rhizosphere and root endosphere act as ‘gatekeepers’ to nonrandomly selection of soil microbes, resulting in phylogenetic conservation within these niches (Liu et al., 2017). The plant recognises these microbes by perceiving their conserved molecular signals- microbe-associated molecular patterns (MAMPs) through the plant high-affinity pattern-recognition receptors on the cell surface. JA and SA are critical hormones in plant immunity, where JA is mainly involved in plant responses to necrotrophic pathogens while SA is involved in the biotrophic and hemibiotrophic pathogens (Bari and Jones, 2009). The two pathways communicate with each other (hormone crosstalk) to orchestrate plant immune responses to pathogens (Bari and Jones, 2009). Recent evidence suggests strong modulatory roles of JA and SA signalling for the controlled colonisation of the commensals in plants (Lebeis et al., 2015). For example, it has been demonstrated that JA and SA signalling mediated the microbiome assembly in the root and rhizosphere of *Arabidopsis* plants (Carvalhais et al., 2013; Lebeis et al., 2015). As a further step to these findings, our study provides evidence that plant microbiomes in return can mediate regulations of plant defences (esp., the SA signalling pathway) when under pathogen attack. This observation can be explained by the fact that (i) the rhizosphere microbiome directly interacts with the plant immune system, for example, they can induce the MAMPs-triggered plant immunity in plants (Teixeira et al., 2021), and (ii) root microbiomes directly interact with the fungal pathogens, and influence disease infection outcomes on wheat (Seneviratne et al., 2007; Hoffman Michele and Arnold, 2010). Overall, our results for the first time demonstrate an emerging role of the plant microbiome in the three-way interaction between the plant, the pathogen and the environment. Amendment of current conceptual frameworks of disease triangle to explicitly consider plant and soil microbiomes will be needed to understand the impacts of fungal pathogens on plant pathogenesis.

### Fp-infection on wheat, pathways and colonisation

The rhizosphere and roots are the first contact point for soil-borne pathogens to invade and for their interactions with other non-pathogenic microbes. We found that *Fp* loads in the rhizosphere and root endosphere (revealed by ITS amplicon sequencing for both cases), base stem (revealed by qPCR analyses) and the plant disease severity (visually rated) all significantly correlated with each other, and the pathogen load was always higher in the diseased plants than healthy plants. This suggests a progressive infection of *Fp* on healthy wheat, where pathogens carried on plant residues in soil first invade the rhizosphere, then colonise and infect the roots, and further proceed to base stems via xylem, causing the typical CR syndrome – stem brown discolorations (Hogg et al., 2007) (Fig.5). Stem discolorations are suggested to be the most reliable indicator of the CR disease at mid to late grain fill stage of wheat without uprooting the plant (Hagerty et al., 2021). However, CR-disease detection based on stem discoloration is often too late to adopt any disease interventions as the plant at this stage has been severely damaged. Our approach of using amplicon sequencing to profile the rhizosphere/root microbiomes was efficient in detecting soil/plant-colonised pathogens, which perhaps can accurately predict CR disease because it detects the pathogen and quantifies its abundance before disease symptoms become obvious. We also detected a small amount of *Fp* (<1.2%) in wheat leaves. *Fp* in the aboveground plant can lead to head infection, namely the *Fusarium* head blight disease (Miedaner et al., 2008). This is because *Fp* produces macroconidia at base stem of CR-infected plants, which can be dispersed up to the canopy by rain-splash (Obanor et al., 2013) or potentially via plant vascular systems. The spores also likely infect heads at wheat flowering stage and cause FHB symptoms (Obanor et al., 2013).

**Fig. 5.**
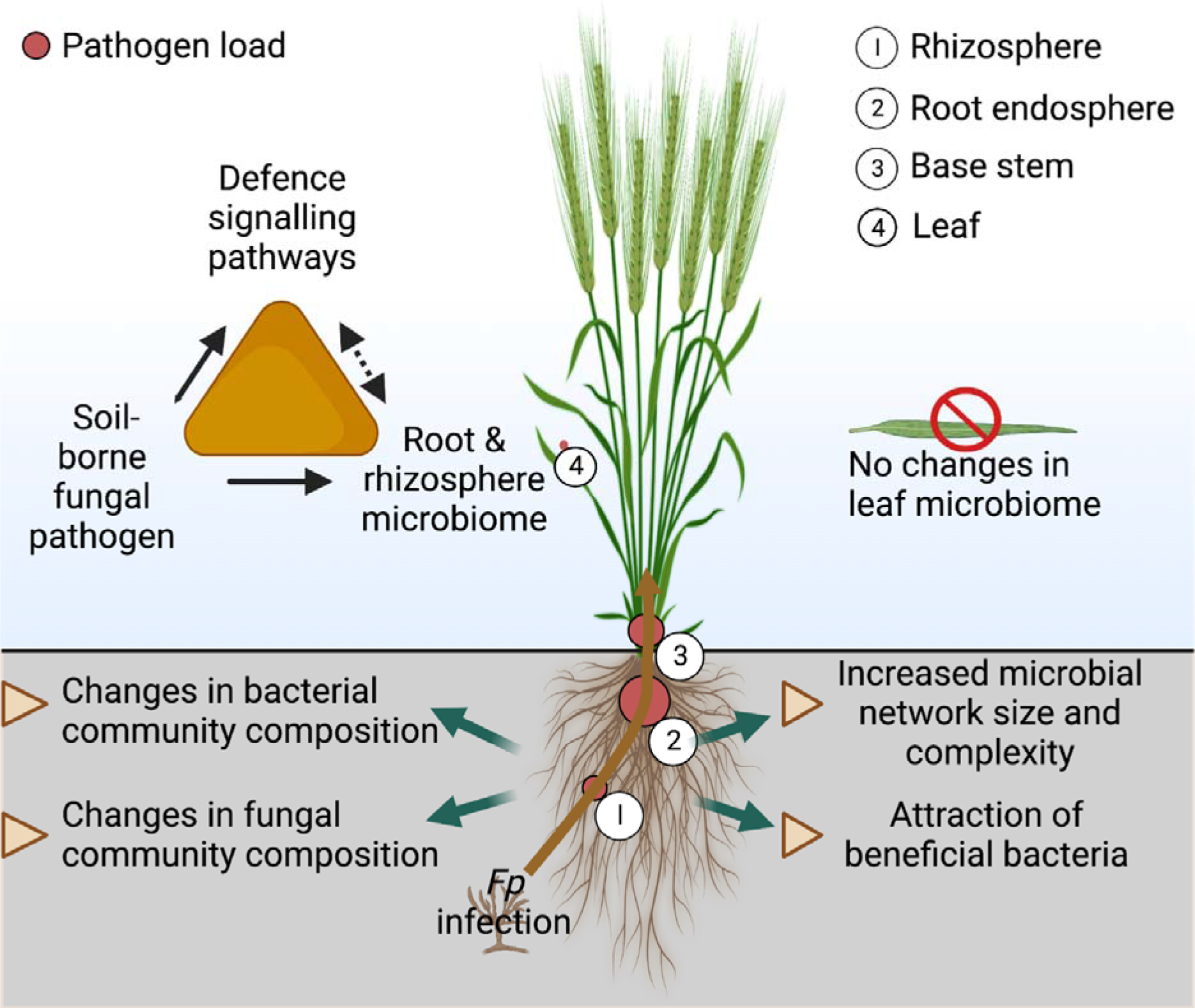
Crown rot (CR) disease infection and induced changes in wheat microbiomes and defence. The level of pathogen (*Fp*) colonisation in fungal communities increased from the bulk soil to the rhizosphere and the root endosphere. The pathogen further transferred to the base stem, where it caused tissue damage and stem discolouration. The pathogen could be detected on the leaf but only presented in a small proportion of the leaf fungal community and did not correlate to the disease infection. Furthermore, wheat defence signalling pathways, wheat microbiomes and the pathogen intimately correlated with each other, and microbial co- occurrence network complexity was higher in *Fp*-infected plants relative to healthy plants. Amendment of current conceptual frameworks of disease triangle to explicitly consider plant and soil microbiomes and their interactions with the plant immunity is needed to understand the impact of fungal pathogens on plant pathogenesis.

### Fp-infection impacts on the wheat microbial composition in the root and rhizosphere but not the leaf environment

The bacterial and fungal community composition in the rhizosphere and root endosphere was influenced by the pathogen infection. This was presumably driven by the direct interactions between the fungal pathogen and soil/plant microbiomes (Liu et al., 2020b), and indirectly by the disease-induced alterations of plant physiology, particularly via root exudates that shape the belowground microbiomes (Yuan et al., 2018). Biological implications of such microbial shifts on wheat performance have been investigated in our previous study, where we found with the presence of the *Stenotrophomonas* sp. in soil promoted wheat growth and disease resistance (Liu et al., 2021), which supports a ‘cry for help’ strategy in wheat plants. In the current study, we also found a few other abundant bacterial OTUs with relative abundances >1% were linearly correlated with plant disease severity, including a *Pseudomonas* sp., a *Rhizobium* sp. and a *Microbacterium* sp. Bacterial strains affiliated to these species also possess plant growth promoting traits, such as ammonia and phytohormone production as well as pathogen inhibition (Liu et al., 2017). Consistently, previous studies showed that the microbial community composition and function of the barley plant (*Hordeum vulgare*) changed upon bacterial or fungal pathogen attack, and microbial traits of disease suppression (e.g., *fluorescent pseudomonads*, genes *phlD* encoding 2,4-DAPG) were enriched in the infected plant rhizosphere (Chapelle et al., 2016; Dudenhöffer et al., 2016; Ginnan et al., 2020). Similarly, plant roots enriched pathogen inhibiting bacteria *Chitinophagaceae* and *Flavobacteriaceae* and functional traits with the presence of pathogens in soil (Carrión et al., 2019). In our study, a *Rhodoferax* sp. in the wheat root endosphere had a significant correlation with the pathogen load, however, host functions provided by this bacterium are unclear. Interestingly, the enriched fungal species, *Sarocladium strictum,* in the rhizosphere has been previously reported as a biocontrol agent against the *Fusarium* head blight (Rojas et al., 2020), indicating that wheat plants also recruit fungal species agents to suppress disease incidence. This result further supports the cry for help strategy of the wheat plant.

### Fp-infection increased the complexity of the root-associated microbial co-occurrence network

Microorganisms live within complex networks in the plant compartments, where extensive microbial interactions occur, such as competition and cooperation among microbes (Hassani et al., 2018). Among these, certain closely interactive microbes can form ecological clusters with co-occurring species sharing common environmental preferences (Delgado-Baquerizo et al., 2018). The diseased wheat plants demonstrated a microbial co-occurrence network with a greater size and higher number of interactions than those in healthy plant microbiomes. This pattern was observed for both the bacterial and fungal communities in both the rhizosphere and root endosphere of the wheat plant, indicating consistent responses of different plant compartments to the disease infection. Consistent results were also reported before, where disease/pathogen infections induced a higher microbial network complexity in the plant rhizosphere and/or root endosphere (Alahmad et al., 2018; Hu et al., 2020). For example, highly connected microbial networks occurred when soil microbiota face environmental perturbation, e.g. inoculation of pathogens in a disease-suppressive soil (Carrión et al., 2019). In this study, network analyses revealed that 80% of the interacting nodes in the pathogen-inoculated suppressive soil belonged to *Chitinophaga*, *Flavobacterium* and *Pseudomonas* species, which possess significant potentials in pathogen inhibition (Carrión et al., 2019). Consistently, in our study, we found that the (potential) beneficial bacterial species such as OTU_89589 (a *Stenotrophomonas* sp.), OTU_85258 (a *Pseudomonas* sp.), OTU_121632 (a *Rhizobium* sp.), OTU_117739 (a *Microbacterium* sp.), and OTU_53413 (a *Rhodoferax* sp.) were well connected in the networks but without dominant roles being found. Overall, our findings provide evidence that disease infection by *Fp* affects the spatial heterogeneity of plant microbiomes that varies between plant compartments (leaf, rhizosphere and root endosphere).

### Functional biomarkers for crown rot disease

The metagenomic analyses detected a relatively large variance of both microbial community and functional gene structure among wheat rhizosphere samples, which is probably due to large variances of field samples in their microbial community structure and functions. However, we were able to detect disease-induced effects on the rhizosphere microbiome and revealed a range of functional genes that differed between samples, which are likely functional biomarkers of the wheat crown rot disease. For example, chorismate mutases (enriched by *Fp*-infection) catalyse the conversion of chorismate to prephenate in microorganisms (Romero et al., 1995). Chorismate is also a precursor for plant defence-related hormones of salicylic acid and indole-3-acetic acid, aromatic amino acids, and other metabolites (Dewick, 1995). Therefore, an increased abundance of this gene in the rhizosphere may have implications in wheat plant growth, development and defence. However, this is only a preliminary finding and it requires future research to investigate in-depth. Those downregulated genes in the rhizosphere also play important roles in plant-microbe interactions. For example, the *gcdH* gene that encodes for a glutaryl-CoA dehydrogenase can be involved in many different plant signalling pathways such as metabolisms of tryptophan and other secondary metabolites (Nouwen et al., 2021). Similarly, these genes are interesting for the future plant-microbe interaction research esp. under disease infections on plants.

## Conclusions

Our results demonstrated that *Fp*-infection of wheat plants altered the rhizosphere and root- colonised microbial communities; while in contrast, effects on the phyllosphere microbiomes were limited. This result suggests that the belowground plant-associated microbial communities play crucial roles in plant responses to pathogen infections on plant. This is supported by our co-occurrence microbial network analyses, where the rhizosphere and root-associated microbial networks showed higher complexity in diseased plants than healthy plants. More importantly, this study revealed a three-way interaction between the plant microbiome, the pathogen and plant defence signalling pathways. This result highlights a critical role of the microbiome in mediating plant defence responses to pathogen attack. Overall, our findings revealed novel understandings of disease-induced microbial changes in wheat, which may accelerate the development of novel crop-optimised microbiome products to sustainably control soil-borne diseases in cereal production industry.

## Author contributions

H.L., J.L. and B.S. conceptualized the idea; H.L., J.W., M.D.B., and H.Z. analysed the data; HL did the writing with all authors having critically revised the manuscript.

## Data accessibility

The 16S rRNA and ITS2 amplicon sequences associated with this study have been deposited in the NCBI SRA under accession: PRJNA436828.

